# Mapping the Single-cell Differentiation Landscape of Osteosarcoma

**DOI:** 10.1101/2023.09.13.555156

**Authors:** Danh D. Truong, Corey Weistuch, Kevin A. Murgas, Prasad Admane, Bridgette L. King, Jes Chauviere Lee, Salah-Eddine Lamhamedi-Cherradi, Jyothishmathi Swaminathan, Najat C. Daw, Nancy Gordon, Vidya Gopalakrishnan, Richard G. Gorlick, Neeta Somaiah, Joseph O. Deasy, Antonios G. Mikos, Allen Tannenbaum, Joseph Ludwig

**Affiliations:** Department of Sarcoma Medical Oncology, The University of Texas MD Anderson Cancer Center, Houston, TX; Department of Medical Physics, Memorial Sloan Kettering Cancer Center, New York, NY; Department of Biomedical Informatics, Stony Brook University, Stony Brook, NY; McCombs Institute, Department of Clinical Cancer Prevention, The University of Texas MD Anderson Cancer Center, Houston, TX, 77030, USA; Division of Pediatrics, The University of Texas MD Anderson Cancer Center, Houston, TX, 77030, USA; Department of Bioengineering, Rice University, Houston, TX; Department of Applied Mathematics and Statistics, Stony Brook University, Stony Brook, NY; Department of Computer Science, Stony Brook University, Stony Brook, NY

## Abstract

The genetic intratumoral heterogeneity observed in human osteosarcomas (OS) poses challenges for drug development and the study of cell fate, plasticity, and differentiation, processes linked to tumor grade, cell metastasis, and survival. To pinpoint errors in OS differentiation, we transcriptionally profiled 31,527 cells from a tissue-engineered model that directs MSCs toward adipogenic and osteoblastic fates. Incorporating pre-existing chondrocyte data, we applied trajectory analysis and non-negative matrix factorization (NMF) to generate the first human mesenchymal differentiation atlas. This ‘roadmap’ served as a reference to delineate the cellular composition of morphologically complex OS tumors and quantify each cell’s lineage commitment. Projecting a bulk RNA-seq OS dataset onto this roadmap unveiled a correlation between a stem-like transcriptomic phenotype and poorer survival outcomes. Our study takes the critical first step in accurately quantifying OS differentiation and lineage, a prerequisite to better understanding global differentiation bottlenecks that might someday be targeted therapeutically.

**Statement of Significance:** OS treatment kills proliferating cells without addressing the root disease cause: dysregulated differentiation. Constructing a roadmap of mesenchymal differentiation enabled us to identify gene modules linked to OS cell fate and patient survival. This ability to characterize and quantify cancer cell fate enables a novel, differentiation-based drug-screening strategy.

## Introduction

Based upon their significant genetic, phenotypic, and lineage-specific diversity, the World Health Organization splits sarcomas into more than fifty unique sarcoma subtypes^1^. Many sarcomas harbor specific chromosomal translocations, oncogenes, or lost tumor suppressors that are used both as diagnostic markers and as potential therapeutic targets^2^. Although genetically heterogeneous, sarcomas are often classified based on their apparent differentiation status and the cell types of the adult mesenchymal lineage that they most resemble. High-grade osteosarcoma (OS), a heterogeneous class of poorly differentiated bone sarcomas, is further subclassified based on predominant tissue features resembling osteoblastic, chondroblastic, fibroblastic, and telangiectatic^3,4^.

With rare exceptions, such as p53-mutated tumors in people with Li-Fraumeni syndrome, most OS have a complex karyotype that is caused by chromothripsis and lack a recurring gene that can be targeted^5^. Although a variety of genetic mutations and genomic structural events (translocations, copy alterations) have been linked to OS, these are often unique to each patient^6^. This presents an obvious challenge in developing therapeutic options broadly applicable to OS patients. Nevertheless, one commonality among high-grade OS is their genetic drift from normal osteoblasts and their varied degrees of dedifferentiation observed along mesenchymal lineages^6^. Therefore, developing tools to accurately identify and quantify the distinct mesenchymal cell populations within OS may better inform patient stratification and therapeutic strategies.

Although single-cell RNA-sequencing (scRNA-seq) has emerged as a promising approach to capture gene expression profiles of individual cells, identifying mesenchymal differentiation states and quantifying stemness still pose unique challenges. Recent tools such as StemID^7^, SCENT^8^, SLICE^9^, and CytoTRACE^10^ have been developed to quantify stemness, but they cannot identify lineage-specific differentiation states. Furthermore, unlike hematopoiesis – which begins during embryogenesis, continues throughout adulthood, and is easily studied with routine blood draws – most connective tissue development occurs during early fetal development and thus hasn’t been readily studied outside of pathological scenarios (e.g., during bone fracture repair). Finally, projecting cancer cells onto a normal differentiation landscape is fundamentally challenging because tumor expression profiles only partially overlap with normal tissue expression profiles.

The present work demonstrates a joint experimental and computational strategy for mapping osteosarcoma onto its underlying mesenchymal differentiation landscape. We developed a high-resolution scRNA-seq reference map composed of osteogenic, adipogenic, and chondrogenic lineages derived from human primary mesenchymal stem cells. Our approach to enrich for differentiating mesenchymal cells utilizes an ex vivo tissue engineered model, adapted from established methods developed by the regenerative medicine field in their attempt to create functional connective tissues coaxed with the addition of lineage-specific growth factors and biomechanical cues^11^. We hypothesized that this Mesenchymal Tissue Landscape (MTL) would catalog the various differentiation states possible in OS.

To quantify differentiation states in the MTL, we applied archetype analysis based on Normalized Nonnegative Matrix Factorization (N-NMF) to identify recurring gene expression signatures that accurately captured lineage-specific, temporal dynamics^12^. These signatures were then applied to estimate the relative compositions of each cell lineage and differentiation state within single-cell OS tumor data, including patient-derived xenograft (PDX) and tumor biopsy samples. Notably, individual OS cells were composed of multiple archetypes, including those that predominantly express genes related to mesenchymal progenitor, osteogenic, and chondrogenic signatures. This suggested that unlike normal cells, which may be reliant on a single archetype, cancer cells co-opt programs from multiple archetypes^8^. Additionally, we observed that the estimated archetypes of each tumor were associated with predominant tissue features (osteoblastic, chondroblastic)^13^. Lastly, we applied our decomposition approach to bulk OS tumor samples, finding that a signature of advanced differentiation was associated with improved survival outcomes.

The ability of our approach to determine lineage and differentiation states in distinct datasets suggests that this approach can be more generally applied to different cancers along the mesenchymal differentiation landscape or possibly even carcinomas where differentiation landscapes remain in their infancy.

## Results

### Intratumoral Heterogeneity in Osteosarcoma PDXs

As proof of concept before exploring human mesenchymal differentiation, we analyzed single-cell transcriptomes from three OS PDXs to explore the heterogeneity of cell types and differentiation states^13,14^. These three PDXs included OS1 (an osteoblastic osteosarcoma), OS31 (an osteosarcoma of unknown subtype), and SA98 (a chondroblastic osteosarcoma) (see Supplemental Table S1 for PDX clinical demographics)^13^. After filtering for high-quality cells by scRNA-seq metrics, we obtained 19,538 OS cells for analysis. Initial visualization of the uniform manifold approximation and projection (UMAP) showed that cells clustered mainly by PDX sample (Fig. 1A). Given the striking degree of intratumoral heterogeneity, we used a Louvain graph-based clustering method to identify 15 distinct clusters (Fig. 1B)^15^. While many clusters are composed of one PDX, some comprise more than one, suggesting the presence of conserved states between different OS PDXs.

**Figure 1:**
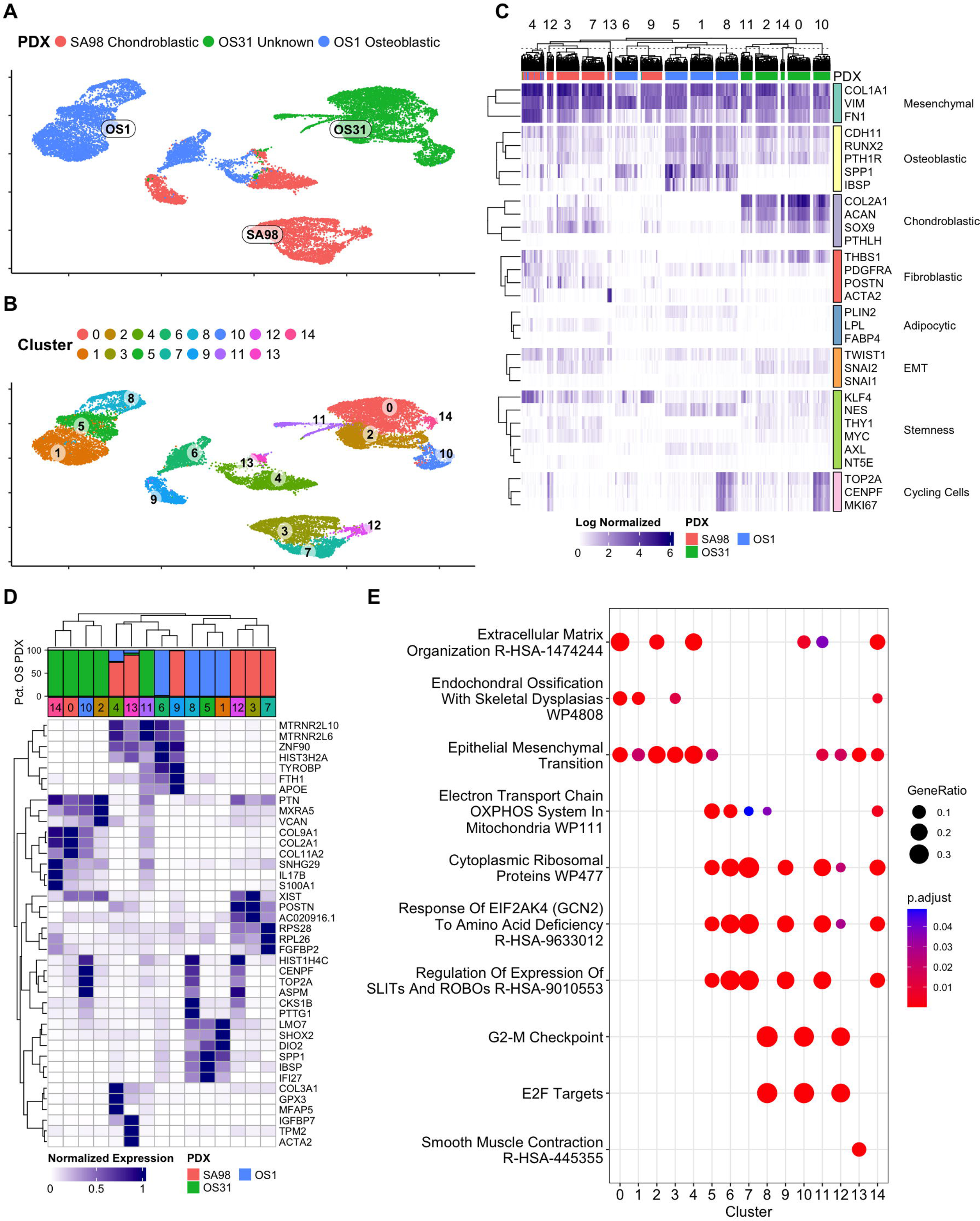
Single-Cell Sequencing Reveals Intratumor Heterogeneity in Patient-Derived Xenograft Models of Osteosarcoma. **A:** Single-cell gene expression UMAP of three OS PDX samples. **B:** Louvain clustering identified 15 clusters of cells in the three PDXs. **C:** Heatmap of gene expression markers (normalized by maximum across all cells) of all individual cells. Genes were grouped according to specific lineage markers or biological processes. **D:** Average cluster gene expression (normalized by maximum across all clusters) of the set of top 3 differentially expressed genes in each cluster. **E:** Dot plot of pathway analysis scores for each cluster.

We then selected classical markers of the cell phenotypes expected in OS PDXs (e.g., osteoblasts, chondroblasts, and fibroblasts), as well markers of adipocytes that served as a negative control (Fig. 1C). Significant heterogeneity in osteoblastic markers among the three PDXs was observed. The OS1 PDX (clusters 1, 5, and 8) displayed the strongest expression of the osteoblastic markers *RUNX2*, *CDH11*, and *SPP1* but also contained a subpopulation of fibroblastic cells and another that lacked strong osteoblastic or fibroblastic markers. The OS31 PDX (clusters 0, 2, 10, 11, 14) was predominantly chondroblastic and had the strongest expression of the chondroblastic markers *COL2A1*, *SOX9*, and *ACAN*. SA98 was not predominantly any one differentiation state but had expression of osteoblastic, chondroblastic, and fibroblastic markers. All PDXs also had a subpopulation of cycling cells, as demonstrated by the expression of *MKI67* and *TOP2A*. We also explored markers of EMT and stemness but did not detect a significant trend in the PDXs, which might explain the expected rarity of cancer stem cells^16^.

Using the clusters, we performed differentially expressed gene (DEG) analysis and ranked the top-expressed markers for each cluster (Fig. 1D; Supplemental Table S2). Clusters 4 and 13 are comprised of mostly SA98 and OS1 PDX cells with fibroblastic genes (*COL3A1*, *FBN1*, *ACTA2*) and lacking osteoblastic genes. In addition, we observed additional evidence of fibroblastic expression for SA98 in Clusters 3, 7, and 12 (*POSTN, FGFR1, FGFBP2*), some evidence of a chondroblastic phenotype (*COL11A1*), and a lack of osteoblastic expression. Additionally, clusters 8, 10, and 12 were indicative of cells undergoing cycling. Enrichment analysis showed four possible major gene expression programs that are active in these three PDXs. Many clusters (0, 2, 3, 4, 10, 14), which were predominantly composed of OS31 and SA98, were associated with ECM-related gene sets (Fig. 1E; Supplemental Table S2). Interestingly, the predominantly osteoblastic OS1 lacked many of the ECM gene sets. Conversely, cluster 1 from OS1 had heightened activity of osteoblast differentiation and bone development genes. Interestingly, cluster 13 was enriched with contractile-related genes, including *TPM2* and *ACTA2*.

### Constructing a Map of the Mesenchymal Transition Landscape

Since such few cells transit mesenchymal linages postnatally, it isn’t yet possible to detect and profile these rare cell types in sufficient numbers using existing technologies in children or adults. Instead, we utilized an *in vitro* model of human mesenchymal differentiation^11^(Fig. 2A). Initially, scRNA-seq was used to profile the mesenchymal differentiation landscape of MSCs under various differentiation conditions and map the changing transcriptome as cells reached their intended terminal fate. To do so, we applied a bioengineered differentiation model involving biochemical and biophysical signals to direct osteogenic and adipogenic tissue differentiation^11^. We then collected cells at defined time points to generate a single-cell time-course profile of each lineage.

**Figure 2:**
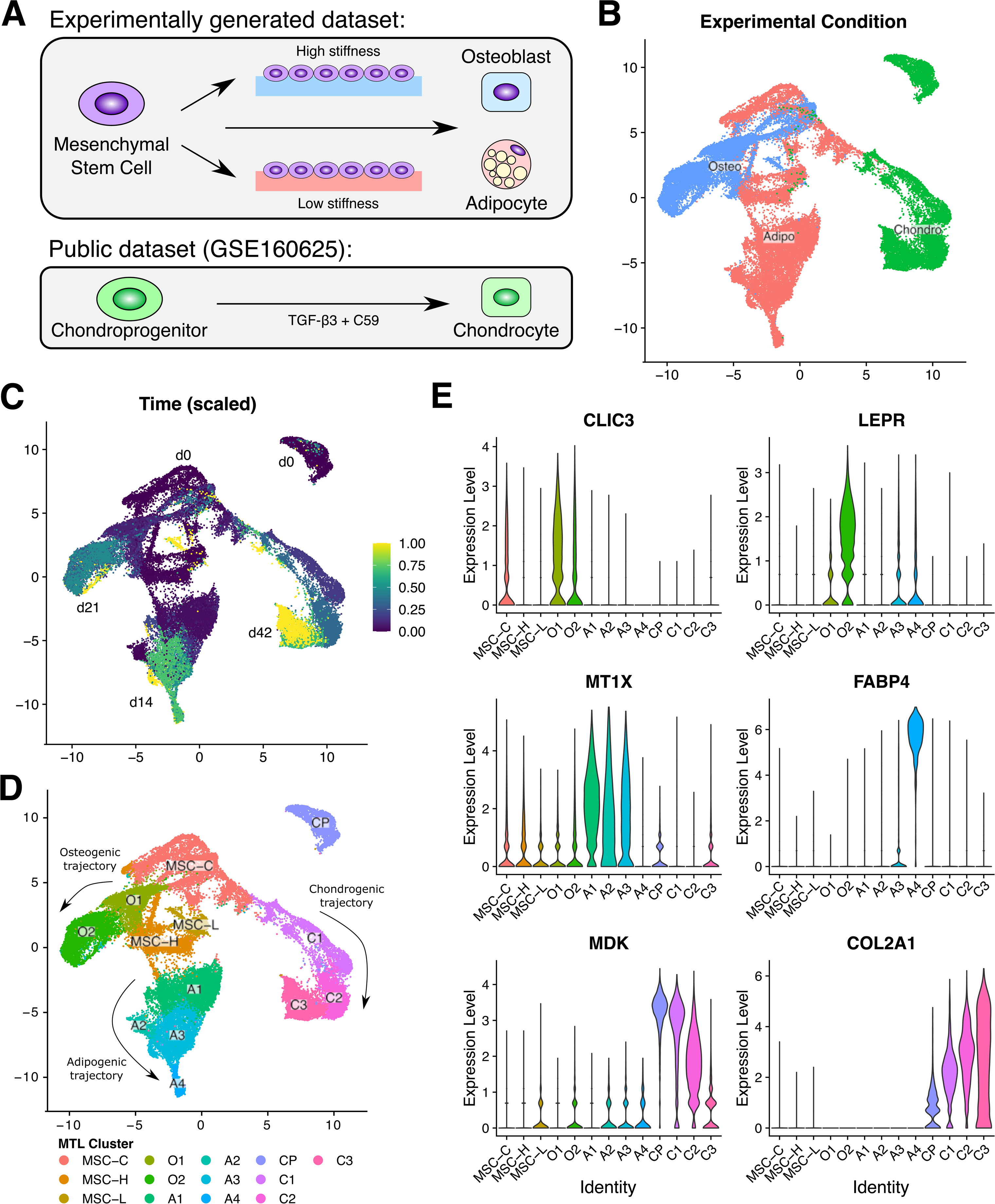
Constructing a Differentiation Atlas of the Mesenchymal Tissue Landscape. **A:** Schematic of datasets utilized to construct an atlas of the Mesenchymal Tissue Landscape (MTL). Osteogenic and adipogenic lineages were experimentally generated over time, culturing on hydrogels of varying stiffness to influence differentiation trajectory. The chondrogenic lineage was sourced from a publicly available dataset (GSE160625), which measured a time course of chondrogenesis in cultured chondroprogenitor cells treated with a combination of TGF-β3 and C59. These two datasets were integrated to construct a mesenchymal differentiation map containing three lineages. **B:** UMAP of integrated MTL colored by experimental condition specifying the differentiation protocol in which the cells were cultured. **C:** UMAP colored by experimental time, scaled to the endpoint of each experiment (d21 for osteo, d14 for adipo, d42 for chondro). **D:** UMAP with designated clusters along three distinct lineages. Clusters were manually annotated as mesenchymal stem cell (MSC), osteogenic (O1-O2), adipogenic (A1-A4), chondroprogenitor (CP), chondrogenic (C1-C3). **E:** Violin plots of marker genes for early and late stages of each lineage showing distinct temporal patterns.

Because our differentiation strategy only modeled osteogenic and adipogenic differentiation, we could not directly estimate the chondrogenic lineage. Chondrogenesis, of course, is an important mesenchymal-origin cell lineage and constitutes a major component of some chondroblastic OS tumors^6,17^. To address this challenge, we augmented our experimentally generated osteogenic and adipogenic MTL with a publicly available chondrogenic lineage single-cell dataset^18^. In brief, the study generated human iPSC-derived chondroblast precursor cells that were then chemically stimulated to undergo chondrogenic differentiation. Throughout their precursor-to-chondrocyte lineage transition, gene expression was assessed by scRNA-seq. With similar time-course data, a batch correction procedure was performed to integrate the chondrogenic lineage with the experimentally generated osteogenic and adipogenic lineages. Subsequently, these data were examined to investigate lineage-specific and temporal gene expression dynamics on the mesenchymal tissue landscape (MTL).

Using the integrated cell embedding produced by Harmony, we generated a Uniform Manifold Approximation and Projection (UMAP) to visualize the topological structure of our single-cell data^19^. Using Louvain graph-based clustering of the Harmony embedding^19^, we generated 13 distinct clusters representative of MSCs, osteoblasts, adipocytes, chondrocytes, and transitional states. We utilized the known condition and time-point data to manually annotate the clusters (Fig. 2B-D)^11^. Our analysis identified two osteo-lineage clusters (O1-O2), four adipo-lineage clusters (A1-A4), three chondro-lineage clusters (C1-C3), in addition to a cluster of cycling MSCs (MSC-C), two populations of MSCs with differing expression of YAP/TAZ-related genes (MSC-H and MSC-L; Supplemental Fig. S1), and a distinct chondroprogenitor cluster (CP). At the earliest time points that occur before the MSCs underwent differentiation, we observed significant overlap between MSCs and the cells exposed to conditions used to instantiate adipogenic or osteogenic lineage commitment. While this implies, expectedly, a shared mesenchymal cell origin, we don’t intend to suggest that osteoblasts and adipocytes share a common lineage; the overlap simply indicates the same MSC cell lines were used to initiate each experiment^20^. Across the clusters, we observed lineage-specific genes with distinct temporal expression profiles corresponding to early and late timepoints (Fig. 2E).

### Gene Markers Validate the MTL Map and Quantify Differentiation along Multiple Mesenchymal Lineages

Cluster-specific differentially expressed genes (DEGs) were identified using pairwise differential expression analysis in Seurat, comparing each cluster against the rest (Supplemental Table S3). The analysis demonstrated consistency between the cluster identities and known lineage-specific markers and pathways (Supplemental Fig. S2). We observed high expression of cell-cycle genes (*CENPF, MKI67, TOP2A*) in MSC-C and CP clusters and to some degree in cluster C1^21^. This was further illustrated by scoring a cell-cycle gene set and identifying the G2M, S, and G1 cell-cycle phases (Supplemental Fig. S3). We observed clear localization of cycling cells in G2 and S phases in the MSC-C, CP, and C1 clusters. Clusters MSC-H and MSC-L, which consisted mostly of MSCs prior to induction, were enriched for MSC-specific markers (*ENG, NT5E, PRRX1, THY1*)^22^. Since MSCs were cultured on gels of low and high stiffnesses (low = 1.5 kPa, high = 25-40 kPa) and account for differences in stiffness conditions, we identified YAP/TAZ-regulated genes (*ANKRD1, CTGF, CYR61, IGFBP5, TEAD1*) enriched in cluster MSC-H versus cluster MSC-L (Supplemental Fig. S1)^23^.

#### Osteogenic Lineage

Within the two osteogenic lineage clusters, cluster O1 consisted primarily of earlier time points (days 1-3) while cluster O2 consisted of later time points (days 5-21). Our analysis showed that cluster O1 up-regulated genes related to both YAP/TAZ signaling and early osteogenesis, such as *ANKRD1, AXL, CTGF, DKK1,* and *HHIP.* Activation of YAP/TAZ signaling has been shown to directly target *ANKRD1, AXL, CTGF,* and *DKK1* when cells are grown at high stiffness prior to osteogenesis^23–25^. In addition, the expression of *HHIP* suggests regulation of Hedgehog signaling. Previous reports stated that YAP activation drove GLI2 nuclear accumulation, which directly activated GLI targets^26,27^. GLI target up-regulation (*CCND1, FOXC2, GLIPR1,* and *IGFBP6*) within cluster O1 of our model acts to validate this finding experimentally. Within clusters O1 and O2, we observed increased expression of ECM genes (*ACAN, COL1A1, COL1A2, COL8A1,* and *ELN*). Furthermore, expressions of *CRYAB, LEPR,* and *TAGLN* indicate positive regulation of osteogenesis^28–30^. *COL1A1* and *COL1A2* are characteristic of osteoprogenitor cells, while *ACAN* was expressed in chondroprogenitor cells. Additionally, many genes (*DCN, FN1, GSN, IGFBP2, IGFBP6*, and *SERPINE2*) of the osteoblast secretome were detected within cluster O2^31^. Cluster O2 was also highly enriched with additional osteoblast secretome-related genes (*CHI3L1, FBLN1, SAA1, THBS2*). *SAA1* expression has been shown to be induced by osteogenic conditions while also promoting osteogenesis and bone mineralization^32^. In addition, IGFPs, including *IGFBP2, IGFBP4,* and *IGFBP7,* and *IGF2* are some of the osteogenic growth factors up-regulated in cluster O2^33^. Expression of transcription factors *FOS* and *CTNNB1* (β-catenin) indicates activation of mechanosensing pathways in osteoblasts^33^. Osteoblast-specific glycoproteins and ECM proteins (*ECM2, GPNMB*) were highly expressed in cluster O2^34–37^. Likewise, WNT1-Inducible-Signaling Pathway Protein 2 (*WISP2*) was highly expressed in cluster O2 and has been shown to both promote osteogenesis and repress adipogenesis^38,39^.

#### Adipogenic Lineage

In the four adipogenic lineages, cluster A1 primarily consisted of cells from early time-points (days 0.5-3), with clusters A2 (days 3-7), A3 (days 7-10), and A4 (days 10-14) containing successively later time points. Clusters A1, A2, and A3 were highly enriched with metallothioneins (MTs: *MT1X, MT1E, MT1M, MT2A*), echoing similar literature results within 24h of adipogenic induction^40^; MTs are expressed in adipose tissue and have been shown to regulate adipogenic differentiation^28,41^. Clusters A1 and A2 were enriched with *WNT5A, PAPPA,* and *FTH1*. *WNT5A* is a part of the non canonical Wnt pathway activated during adipogenesis^42^. Both *PAPPA* and *FTH1* are up-regulated during activation of peroxisome proliferator-activated receptors (PPARs)^43^. Cluster A3 appeared to be heavily enriched with extracellular matrix (ECM) genes (*COL3A1, COL6A1, COMP, DCN, LUM*). Though not initially enriched during the early phase of adipogenesis, these genes appear to peak within cluster A3. Likewise, various groups have observed that adipogenesis follows a biphasic pattern concerning ECM proteins, where collagens, laminins, biglycan, and lumican demonstrate a decrease upon adipogenic induction but return to basal levels after a certain time, followed by a phase of cell growth and fat storage^44^. As such, cluster A4 was highly enriched for mature adipocyte gene signatures (*ACACB, ADIPOQ, APOE, FABP4, G0S2, FABP5, LPL, PLIN4, PLIN1*).

#### Chondrogenic Lineage

Within the chondrogenic differentiation time course, cluster C1 consisted of primarily day 7, cluster C2 consisted mostly of day 14, and cluster C3 consisted of days 28 and 42. Clusters C1, C2, and C3 all showed increased expression of chondrogenic markers (*COL9A1, MATN4, SOX9*)^18,45^. All three chondrogenic clusters also expressed frizzled-related proteins (*EPYC, FRZB, LECT1*)^46^. Clusters C1 and C2, containing earlier time points, exhibited a few early markers of chondrogenic differentiation (*SOX2, SOX6*), while clusters C2 and C3, containing later time points, exhibited increased expression of additional chondrogenic markers (*ACAN, COL2A1, COL3A1*)^47^. The CP cluster was observed to show distinct transcriptional signatures compared to the rest of the mesenchymal landscape, with expression of genes indicating chondrogenic potential (*SOX2, SOX4*) and some neural crest markers (*FOXD3, PAX3, PAX6, OTX2*) and WNT signaling genes (*MAPK10, WNT4*) reflecting the derived progenitor status of this cluster^48,49^.

### Archetype Analysis Defines Lineage-Specific and Time-Dependent Expression Profiles within the MTL

Next, we sought to quantitate time-dependent signatures of mesenchymal stem cell differentiation along the MTL-derived lineages. We applied archetype analysis to identify temporal and lineage-specific gene expression signatures and subsequently estimate the relative contribution of each archetype in individual cells^50^. Twelve distinct signatures (Supplemental Fig. S4) captured gene expression variability as cells differentiated along the MTL (Fig. 3A). Each cluster in the MTL was characterized by a distinct composition of archetypes, defined based on the highest expressed archetype of each cell (Fig. 3B). More explicitly, the estimated abundance of each archetype reached its peak either at a distinct time or within a particular lineage (Fig. 3C). Archetypes 1-2, for example, captured genes associated with undifferentiated MSCs, whereas archetypes 3-4 were indicative of cells that had begun or completed connective tissue differentiation along adipogenic (red) or osteoblastic (blue) trajectories. Archetype 5 corresponded to an early adipogenic state; by contrast, archetypes 6 and 7 indicated later commitment to their respective adipocyte and osteoblast trajectories. The chondrogenic lineage (green), perhaps explained by the distinct experimental design involved in their profiling, was associated with archetypes 8-12, each peaking at a distinct experimental time. We note the discontinuity between the CP and C1-3 clusters, which we surmise is an artifact observed chiefly due to the sparse 7-day time gap in the external dataset, which failed to capture a rapid cell-state transition between the CP and C1 phenotypes. Overall, the archetypes each reflected a distinct attribute of the MTL, with later clusters expressing both the differentiated and lineage-specific archetypes.

**Figure 3:**
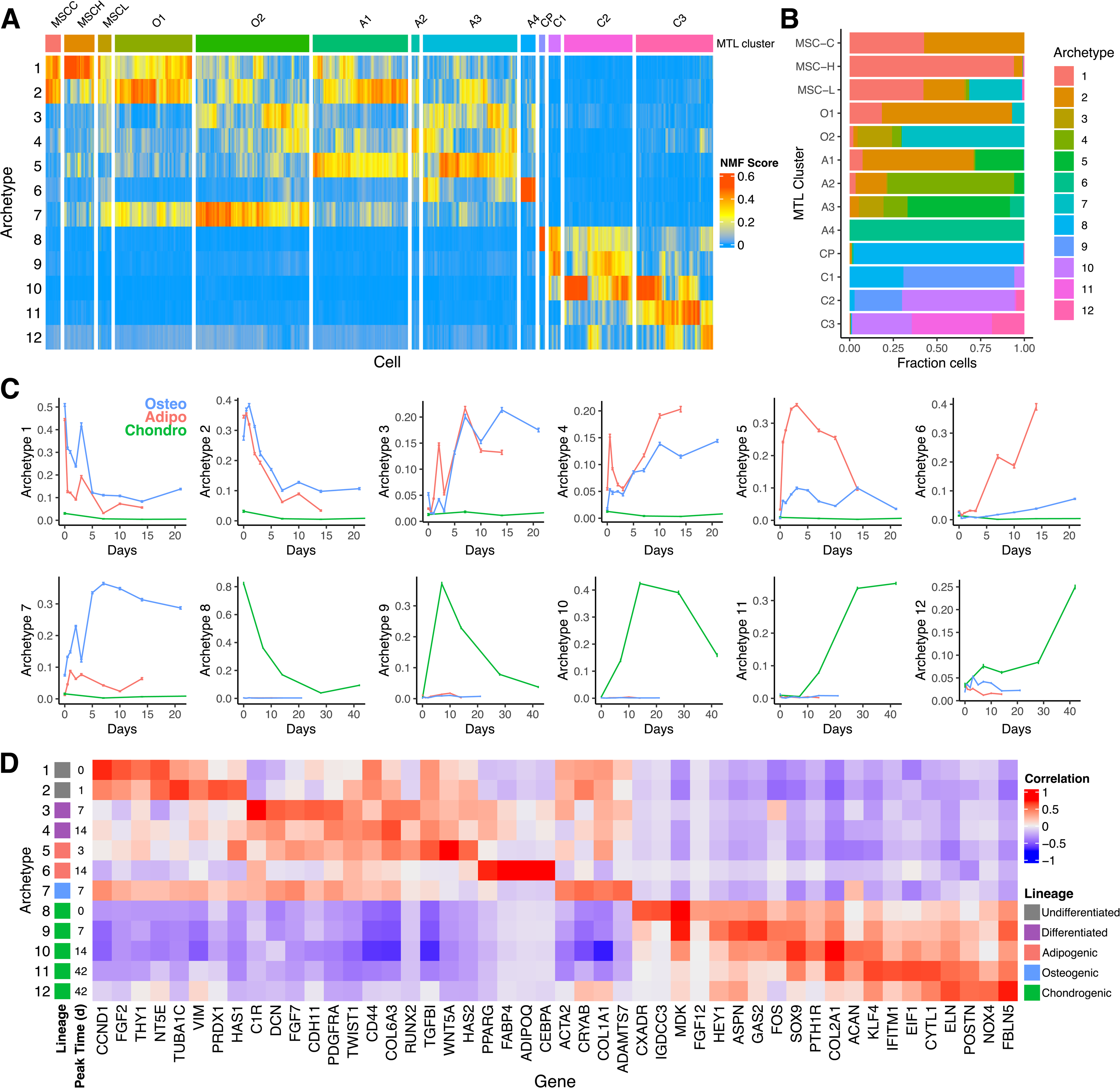
Archetype Analysis Defines Signatures of Distinct Mesenchymal Differentiation States. **A:** Heatmap of archetype score estimates split by MTL cluster. Archetype scores were estimated by normalized nonnegative matrix factorization. **B:** Compositions of each MTL cluster based on each cell’s highest expressed archetype signature. **C:** Average archetype time courses stratified by cell lineage. **D:** Heatmap of representative gene-archetype correlations. Four representative genes were selected among the top Pearson correlates of each archetype based on their known biological relevance. The complete list of correlates is provided in Supplemental Table S4. On left, each archetype was annotated by lineage(s) and peak time.

To confirm that these archetypes capture established lineage-specific and stemness-associated features of gene expression, we conducted a genome-wide Pearson correlational analysis between the expression levels of each gene in our dataset and the single-cell scores assigned to each archetype (Supplemental Table S4, summarized in Fig. 3D). Archetype 1 showed a strong association with well-known MSC genes such *as NT5E* and *THY1*, confirming its representation of an early undifferentiated MSC signature. Archetypes 3 and 4 appeared related to a mesenchymal state, given their correlation to *PDGFRA*, *CD44,* and *TWIST1.* Archetypes 3 and 5 correlated with *RUNX2*, a master regulator of osteogenic differentiation, whereas archetype 6 correlated with *PPARG*, a master regulator of adipogenic differentiation and an antagonist of *RUNX2*. Thus, our results suggest that while *RUNX2* plays a role in both lineages, differentiated adipocytes repress osteogenic gene expression modules as indicated by their expression of archetype 6 and decrease in archetypes 3 and 5. Further evidence that the archetypes captured lineage-specific expression modules, archetypes 6, 7, and 10, were associated with adipogenic (*FABP4*), osteogenic (*COL1A1)*, and chondrogenic (*ACAN)* marker genes, respectively.

### Human Osteosarcoma Tumors Display Distinct Cell State Compositions

To understand osteosarcoma heterogeneity in the context of mesenchymal stem cell differentiation, we projected single-cell expression data from human osteosarcoma PDX (Fig. 4A) and patient (Fig. 4B) samples onto the 12 archetypes defined by the MTL. In contrast to MSC-derived differentiating cells with specific signatures, individual OS cells were comprised of a mixture of several archetypes. To quantify the composition of each tumor, we again classified cells based on their highest expressed archetype (Figs. 4C,D). OS PDX and patient samples exhibited similar archetype profiles, with the major exception being higher expression of archetype 4 (indicative of lineage-indifferent differentiation) in PDX samples compared to patient samples. Notably, archetypes 2 and 3 were predominant in both the PDX and patient samples, indicating activation of stemness and mesenchymal progenitor genes. Strikingly, the high-grade OS tumors analyzed in our dataset harbored only a limited number of cells expressing archetype 7, suggesting impaired osteogenic differentiation. In contrast to osteoblastic OS, chondroblastic OS samples and the PDX of an unknown subtype (OS31) contained cells characterized by the chondrogenic archetype 10, aligning with their classification. In line with prior observations in OS tumor samples (specifically, OS cell group 1)^51^, all three PDX samples and most patient samples contained cell populations predominantly characterized by the adipocyte-associated archetype 6. Importantly, the presence of these populations in the comparatively pure PDX samples signifies a recurring use of adipogenic gene modules, without necessarily undergoing adipogenesis (Supplemental Table S4). Lastly, correlations between pairs of archetypes, like archetypes 2 and 6 as well as archetypes 3 and 10, were consistently observed within the samples and may suggest a conserved mechanism of cell state dysregulation in OS.

**Figure 4:**
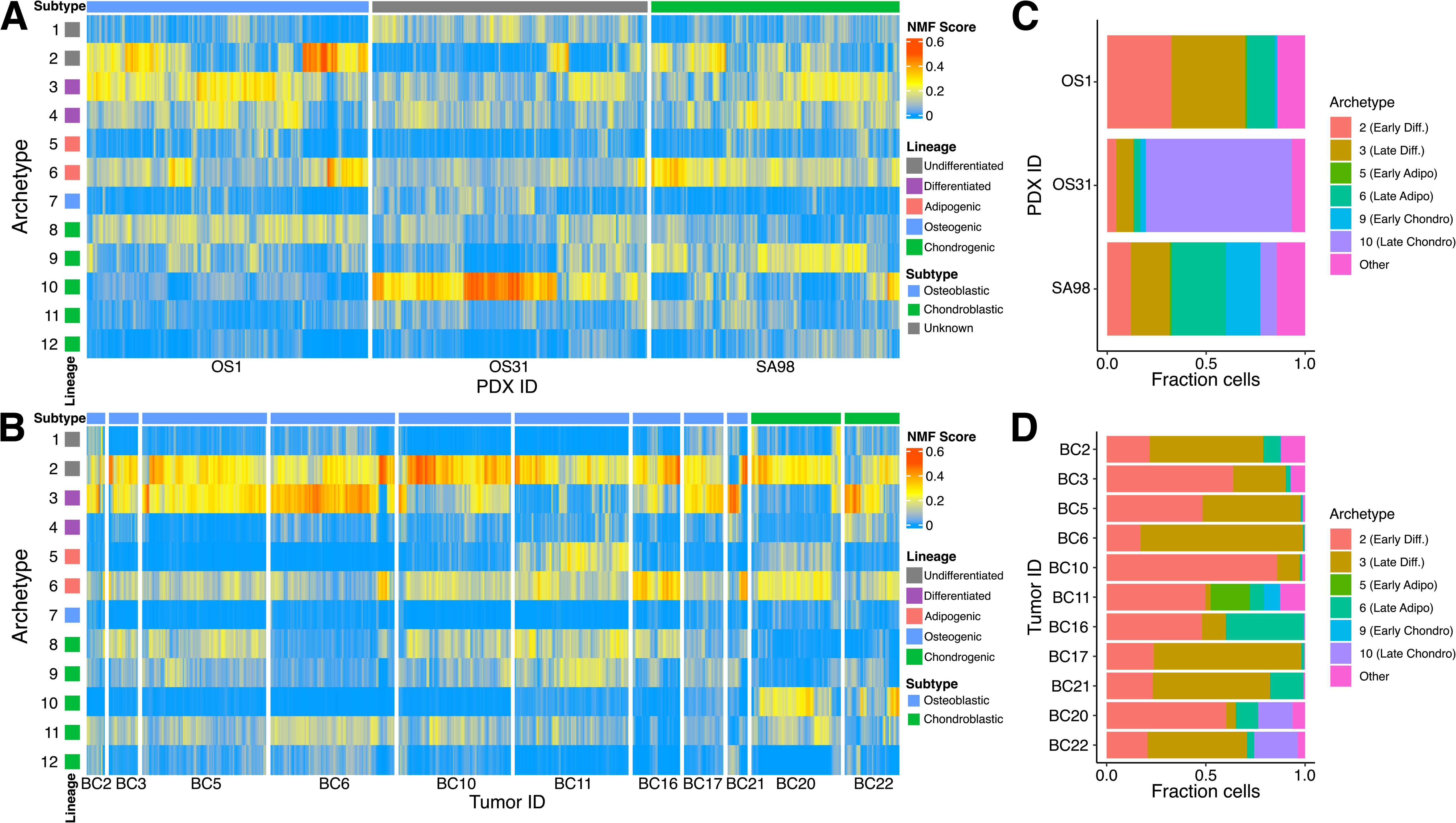
Archetype composition of osteosarcoma tumor samples and PDX models. **A:** Single-cell archetype score heatmap of 3 PDX osteosarcoma models, with hierarchical clustering to accentuate groups of similar cells (dendrogram not shown). Row annotation on the left indicates the lineage of each archetype (same as Fig. 3D). Column annotation indicates the pathologist subtype label based on the predominant cell type. **B:** Single-cell archetype score heatmap of 11 human osteosarcoma tumor samples. Similar annotation as panel A. **C:** Compositions of each PDX based on the maximum archetype score of each cell. **D:** Compositions of each OS tumor.

### A Differentiation Archetype is Associated with Improved Survival in Pediatric Osteosarcoma

We next explored how OS tumor heterogeneity affects clinical outcomes using the NCI’s Therapeutically Applicable Research to Generate Effective Treatments (TARGET) OS data subset, made publicly available to accelerate cancer research. Specifically, we projected the bulk RNA-seq samples from TARGET-OS with overall survival data (N=85) onto the MTL to identify archetypes associated with differential survival outcomes (Fig. 5A). Despite lacking the intratumoral resolution of single-cell analysis, we found substantial variation in archetype compositions across the cohort. Survival analysis in the cohort revealed that gene expression archetype heterogeneity could explain differential osteosarcoma survival (log-rank test: p=0.033; Fig. 5B). A multivariate analysis determined that archetype 3 (a signature of differentiation) was most significantly associated with improved overall survival (Cox PH: p=0.041; Fig. 5C). In other words, a well-differentiated expression signature was associated with improved prognosis as shown previously by many groups^52–55^. Notably, archetype 10, a chondrogenic archetype, was high in a subset of samples, suggesting potential chondroblastic osteosarcoma although subtype information was not available.

**Figure 5:**
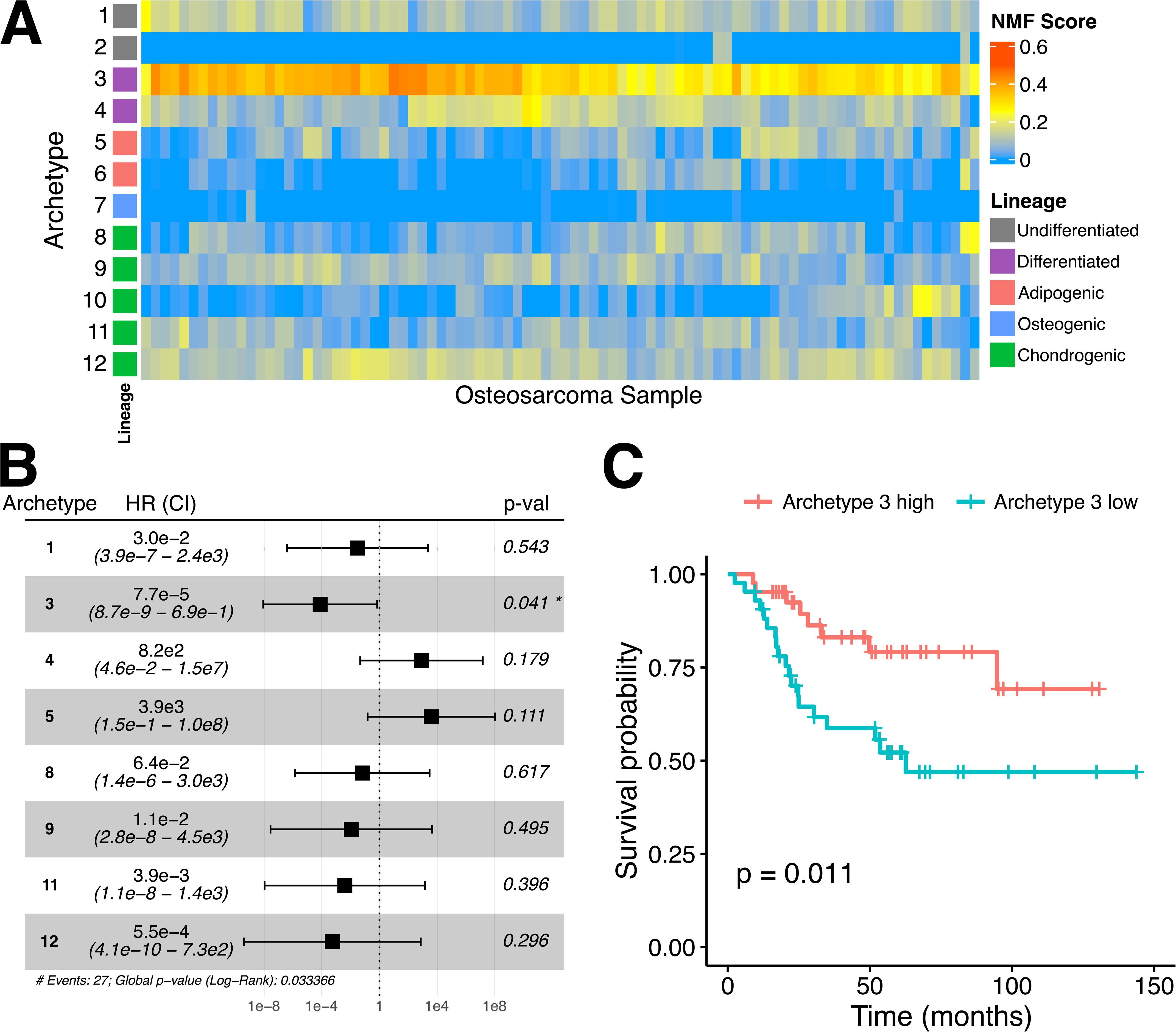
Well-differentiated signature is associated with improved survival in osteosarcoma. **A:** Heatmap of archetype expression in TARGET-OS (N=88); each column is a different patient’s osteosarcoma bulk sample. **B:** Forest plot of the associated Cox regression model with reported hazard ratio (HR) and confidence interval (CI) using the expressed MTL archetypes as regressors. Significance was determined by a multivariable Cox PH test (N=85). **C:** Kaplan-Meier plot stratified by the sample-specific Archetype 3 (differentiated) scores grouped as higher or lower than the median. Significance was determined by a univariable Cox PH test (N=85).

## Discussion

Conventional osteosarcomas appear as spindle-like cells that produce osteoid matrix and are believed to have originated from a malignant transformation of a mesenchymal progenitor. Even though most osteosarcomas are comprised of multiple lineages, they are generally classified as osteoblastic, chondroblastic, or fibroblastic based on their predominant histological features. Without the tools to quantify the lineage of the OS tumor cell composition with a high degree of precision by traditional IHC-based morphometric methods, most high-grade OSs are treated identically using the same cytotoxic chemotherapy regimens developed more than five decades ago. As more powerful proteogenomic methods are applied to tumor samples, one expects to identify prognostic therapeutic biomarkers that ultimately help enable precision-guided medicine that accounts for each patient’s unique tumor characteristics. Here, we take an initial step toward that long-term goal using advanced computational methods that allow one to quantify the differentiation state and lineage of individual osteosarcoma cells. Low-grade OS variants exist that better resemble their mature connective tissue counterparts but are exceedingly rare and were not available, which was a limitation of our analysis.

To better understand osteosarcoma heterogeneity and dysregulated cell differentiation, we developed a Mesenchymal Tissue Landscape (MTL), the first high-resolution tissue-engineered reference map able to profile MSCs by scRNA-seq on their physiological journey toward an osteogenic, adipogenic, and chondrogenic cell fate. Insights generated from the MTL proved valuable in describing mesenchymal differentiation states, revealing distinct lineage-specific gene markers with temporal ordering. It is important to note that we did not observe an osteochondroprogenitor cell in our data, which would represent a shared precursor for osteoblasts and chondrocytes^47^. There are two possible explanations for this limitation. One is that the external data was derived from an iPSC line and not an MSC, like in the data we generated. The second explanation is that the temporal resolution of the chondrogenic data may not have been fine enough and that additional time points at early differentiation are required to observe a shared osteochondral precursor cell. Despite this disconnect, we are still able to learn the osteogenic and chondrogenic differentiation states following this precursor cell. Future studies may help identify the branchpoint where osteoblastic and chondrogenic differentiation diverges by characterizing cells at earlier time points along the osteochondroprogenitor trajectory.

Crucially, the MTL provides a framework to understand OS gene expression patterns in terms of the various mesenchymal lineages. Distinct tumor cell populations expressing stem-like, osteoblastic, adipogenic, and chondroblastic signatures were observed across a representative high-grade OS cohort. Although the adipogenic lineage may not apply to OS, this transcriptomic characterization may prove useful in future studies aimed, for example, at unraveling the drivers of well-differentiated and dedifferentiated liposarcomas, common sarcoma subtypes that lack effective biologically targeted therapies. Further, though the tissue engineering field has demonstrated the ability to push MSCs toward a myoblast phenotype ex vivo by regulating mechanotransduction optimally at 10-12 kPa, this lineage was not included in our initial version of the scRNA-seq MTL, another direction for future studies, in particular for studying defects in rhabdomyosarcoma cell fate^11,56^.

The present study utilized archetype analysis to project osteosarcomas onto the distinct transcriptomic states present in the MTL. Joint analysis of single-cell transcriptomes from three OS PDX models and 11 publicly available human OS tumors enabled us to characterize the predominant osteosarcoma cell states in the data. Given the potential association between these archetype signatures and patient outcomes, this approach could aid in developing patient-specific therapeutic strategies for OS. More OS samples will be required to validate our preliminary findings, potentially enabled through collaboration across the sarcoma research community. Rapid technological changes in the ability to study archival frozen and formalin-fixed paraffin-embedded (FFPE) tissues at the single-cell level, paired with the ability to perform sample multiplexing economically, are on the cusp of enabling these kinds of future large-scale studies.

Methods used to infer cell lineage trajectory or plasticity, such as Waddington-OMT and CytoTrace, have shown tremendous promise in understanding temporal gene expression dynamics^10^. As with any nonlinear method, such results may be challenging to interpret and calibrate. Considering the complex constraints of these methods, such data might be more easily understood by utilizing archetype analysis to define low-dimensional representations of lineage-specific gene expression programs. Here, since the archetypes were trained based on normal mesenchymal differentiation time courses, temporal information derived from archetype analysis could approximate the current differentiation state of sarcoma cells with respect to normal differentiation. Given that many sarcomas may undergo incomplete differentiation or dedifferentiation, this analysis approach provides a way to quantify tumor subpopulations not only of different mesenchymal lineages but also of different temporal stages.

In conclusion, our study has generated a unified and detailed map of the cellular differentiation landscape of three human connective tissue types and demonstrated its potential for characterizing intra-and inter-tumoral OS heterogeneity. Our first-generation MTL lays the groundwork for creating a comprehensive atlas of normal and cancer cell differentiation, which will aid in developing lineage-specific targeted therapies for OS. More broadly, the ability to precisely quantify lineage commitment has far-reaching implications for both basic biology and cancer treatment, like clarifying key genes involved in differentiation and fostering a mechanistic understanding of how tumors dynamically adapt their cell states to evade targeted therapies.

## Methods

### Osteosarcoma Patient-Derived Xenografts

Single-cell RNA-seq data from PDX lines were obtained from a previous study GSE200529^13^. The OS PDXs were isolated upon reaching 2000 mm^3^ and underwent rapid dissociation. The clinical demographics of the PDXs are reported in Supplemental Table S1. The OS1 and OS31 PDXs were obtained from the Houghton lab^14^. The OS1 PDX was derived from a male patient diagnosed with osteoblastic OS who was treatment naïve. The OS31 PDX was derived from a female patient with an unknown subtype and treatment. The SA98 PDX (full ID: MDA-SA98-TIS02) was generated at MD Anderson Cancer Center from a female patient diagnosed with chondroblastic OS having received neoadjuvant chemotherapy^13^. For this study, we selected only “warm” dissociated single-cell RNA-seq data. Each of OS1, OS31, and SA98 are PDX lines maintained by the Pediatric Solid Tumors Comprehensive Data Resource Core^13^.

### Mesenchymal stem cell culture and differentiation

#### MSC isolation and culture

Human primary bone marrow-derived MSCs were isolated from a healthy 23-year-old female donor and provided through an IRB-approved partnership with Texas A&M Institute for Regenerative Medicine. MSCs were selected based on positivity for CD90, CD105, CD73 and negativity for CD34, CD19, CD11b, CD45, CD79a, CD14, HLA-II:DR DQ DP.

#### Polyacrylamide-based hydrogel preparation

Briefly, MSCs were differentiated on polyacrylamide-based hydrogels coated with collagen according to established protocols^11,57^. As *in vitro* differentiation can be inefficient, our differentiation model incorporates biochemical and biophysical cues to better regulate MSC differentiation as described in other works^11,56^. As previously described^11^, we prepare polyacrylamide gels adhered to treated glass slides by glutaraldehyde. The ratio of acrylamide/bis was adjusted for the required stiffnesses. Collagen (10µg/mL) was functionalized to the gels using Sulfo-SANPAH for cell attachment. MSCs were cultured on polyacrylamide-based hydrogels of different stiffnesses (low = 1.5 kPa, high = 25-40 kPa) corresponding to the stiffnesses of the related mesenchymal tissues (adipose ∼ 1-2 kPa^56,58,59^, osteoid ∼ 25-40 kPa^11,60^). Upon adding the differentiation induction medium, created as previously described^11^, cells were collected at specified time points until terminal differentiation.

#### Single-cell suspension, library preparation, and sequencing

Cells undergoing differentiation were collected at specified time-points for sequencing (time points including days 0, 0.5, 1, 2, 3, 5, 7, 10, 14, 21). The cells were washed in PBS before adding trypsin/EDTA (0.25%/380.0 mg/L) followed by 30 min incubation at 37 °C. Afterwards, cells were agitated through pipetting and trypsin activity was briefly terminated by addition of culture media containing serum. Cells were centrifuged and washed with PBS + 0.4% BSA to remove debris and dead cells. Prior to submission for sequencing, cells were re-suspended at 1000 cells/µl.

Single-cell capture, lysis, and library preparation were performed using the Chromium Controller system and established protocols (10x Genomics). Libraries were prepared using Single Cell 3’ v3 with a capture of 1700 cells as input for an estimated target cell capture of 1000 cells due to cell capture efficiency of 65%. Libraries were sequenced using the Illumina NextSeq 500 to sequence each cell at an estimated 50,000 reads per cell. The total number of cells captured was approximately 31,527 cells across all time points with a median unique molecular count of 3.6E4 ± 2.2E4 (SD) per cell and median unique feature of 5.3E3 ± 1.4E3 per cell.

##### Public datasets

###### Chondrogenic Differentiation

Data was accessed at NCBI GEO accession code GSE160625^18^. Raw read counts from each condition were loaded using Seurat R package Read10X command and subsequently merged into a single Seurat data file with added metadata columns specifying the experimental condition and time of each cell. To utilize the authors-reported higher efficiency chondrogenic differentiation with C59 treatment, the chondroprogenitor (Cp) and C59-treated conditions (C59_D7, C59_D14, C59_D28, C59_D42) were selected for analysis^18^.

###### Osteosarcoma Tumors

Data was accessed at NCBI GEO accession code GSE152048^51^. Clinical characteristics of the 11 osteosarcoma tumors including pathological classification are described in supplemental information of the original study. Raw read counts from each condition were loaded using Seurat R package Read10X command and subsequently merged into a single Seurat data file with added metadata column specifying the tumor identity.

###### Pediatric Osteosarcoma (TARGET-OS)

The results published here are in part based upon data generated by the Therapeutically Applicable Research to Generate Effective Treatments (https://www.cancer.gov/ccg/research/genome-sequencing/target) initiative, phs000218. See section “Pediatric Sarcoma Survival Analysis” below for additional details.

### Preprocessing of 10x scRNA-seq data

Cell Ranger (10x Genomics) was used to perform demultiplexing, alignment, filtering, barcode counting, UMI counting, and aggregate the outputs from multiple libraries. Following the standard Cell Ranger pipeline, the filtered gene barcoded matrices were read by Seurat v3 for data preprocessing and initial analysis^61,62^.

As quality control for all datasets, cells with fewer than 500 detected genes were removed, along with genes detected in fewer than 3 cells. Additional outliers of mapped mitochondrial reads and expressed genes per cells were adaptively filtered using the scuttle R package isOutlier command with default threshold of 3 mean absolute deviations. Finally, an absolute threshold of cells with low unique counts (n<1000) and high mitochondrial genes (n>25%) was applied as a filter.

After the removal of low-quality cells, we normalized the data using Seurat R package SCTransform normalization^63^. In brief, SCTransform performs normalization and variance stabilization using regularized negative binomial regression to remove technical variation, returning corrected read counts. Normalization was performed in a batch-specific manner, where the Seurat object of a given dataset was first split by batch or tumor ID, using SplitObject to create a list of separate Seurat objects. SCTransform command with parameters: return.only.var.genes=F, vst.flavor=”v2” was applied to each list object. The resulting normalized data were merged, including the union of genes reported by SCTransform for each batch.

### Harmony Batch Correction

To generate a comprehensive visualization of the MTL, we applied Harmony integration to correct for batch-specific variation among the differentiation data^64^. In brief, Harmony utilizes an iterative correction of the principal components (PCs) to remove batch-specific effects and maximize cell clustering. The data was first normalized using sctransform and a principal-component analysis (PCA) was computed using the top variable genes. The PCA embedding was then batch-corrected by the Harmony algorithm to produce an integrated dimensional reduction with batch correction across tumor_id^19^. Harmony integration was implemented using harmony R package RunHarmony command with parameters: group.by.vars=’Batch.ID’, theta=3, lambda=0.5, tau=300.

### Visualization and clustering

To visualize the mesenchymal landscape, we produced a 2D map by Uniform Manifold Approximation and Projection (UMAP) on the Harmony reduction, storing the reduction as ‘umap_harmony’^65^. Then, using the Harmony reduction, we generated a shared nearest neighbor graph and identified clusters of cells using Louvain algorithm^15^. Resulting clusters were manually labeled based on the predominant experimental condition and time points within each cluster, which largely aligned with three mesenchymal differentiation lineages (osteogenic, adipogenic, chondrogenic) and time-course progression. Heatmaps were produced using the R packages ComplexHeatmap and ggplot2^66,67^.

### Marker Gene Analysis

Differentially expressed marker genes among cell clusters were identified using the FindAllMarkers command of the Seurat R package with parameters: only.pos=T. This method applies the default Wilcoxon rank sum statistical test to perform comparison of all genes in each cluster compared to all other clusters, only considering genes with positive expression and log fold-change (logfc) above the specified threshold and returning a Bonferroni multiple-comparison-adjusted p-value (p_val_adj). Significant marker genes with p_val_adj<0.05 were subsequently assessed to determine cluster-and lineage-specific markers and confirm the expected biological identity of the cluster cell populations. For each cluster, Gene Ontology (GO) pathway enrichment analysis of marker genes was conducted using the enrichGO command of the clusterProfiler R package with parameters: ont=’ALL’, pvalueCutoff=0.01, qvalueCutoff=0.05^68,69^. Marker genes and full enrichment results for each cluster are included in Supplemental Table S3.

### Gene Expression Archetype Analysis

We applied a Normalized Non-negative Matrix Factorization (N-NMF) algorithm to define transcriptional programs or “archetypes” which capture the variability of gene expression patterns across the dataset in a low-dimensional subspace^12^. This approach is a semi-supervised machine learning approach, where we first trained N-NMF archetype coefficients on the mesenchymal differentiation (MTL) dataset and later used these trained archetypes to score the osteosarcoma PDX and tumor datasets. Prior to N-NMF, we filtered out cycling cells (categorized as G2/M or S phase by Seurat’s CellCycleScoring test, keeping only G1 phase cells) to remove the influence of the cell cycle signature. We applied quantile normalization to ensure gene expression matched a similar distribution across all cells. To minimize the impact of sequencing noise present in lowly expressed genes, we isolated differentially expressed genes across the data by removing genes with variance less than 0.70 (corresponding to the bulk of the genes), selecting a total of 454 highly expressed and variable genes across the data. Then, the optimal N-NMF rank was determined heuristically by examining the eigenvalue spectrum elbow plot (Supplemental Fig. S4). N-NMF archetype coefficients were computed using an iterative algorithm to compute a gene-archetype coefficient matrix and cell-archetype score matrix (normalized to sum to 1 for each cell)^70^. Scoring of OS (PDX and tumor) cells with trained archetypes was then computed using a similar iterative algorithm but with the gene-archetype coefficient matrix held fixed as determined in the MTL dataset. Final archetype gene weights and gene-archetype correlations within each dataset are included in Supplemental Table S4.

### Pediatric Sarcoma Survival Analysis

Pediatric sarcoma gene expression profiles were obtained from the Therapeutically Applicable Research to Generate Effective Treatments (TARGET) study OS dataset (TARGET-OS, N=88)^71^. Gene expression data along with clinical phenotype and overall survival data were accessed from UCSC Xena browser, URL: https://xenabrowser.net/datapages/?cohort=GDC%20TARGET-OS^72^. RNA-seq STAR counts were provided as log_2_(CPM+1). For archetype analysis, after mapping gene ensembl_ids to HGNC gene symbol, expression data was quantile normalized with respect to the MTL as the target distribution. Archetype analysis was then used to estimate the composition of archetype gene signatures (trained on the MTL as described above) in each OS. Survival analysis was performed using the resulting archetype score. Initially, low-expressed archetypes which had means across all samples less than 0.05 were filtered out (4 of 12 archetypes were removed). Finally, Cox multiple regression was used to determine the overall hazard ratio and statistical significance of each archetype, as well as the global log-rank statistic.

## Code and Data Availability

All relevant code used for data processing and analysis are available in a public GitHub repository at the following link: https://github.com/Ludwig-Laboratory/sarcoma_differentiation_landscape. The datasets generated analyzed in this study will be available at the Gene Expression Omnibus (GEO, https://www.ncbi.nlm.nih.gov/geo/) upon publication^73^.

## Ethics Statement

The MSCs used in this study were taken from healthy donors through an IRB-approved partnership with Texas A&M. All experiments were conducted per protocols and conditions approved by the University of Texas MD Anderson Cancer Center (MDACC; Houston, TX) Institutional Animal Care and Use Committee (eACUF Protocols #00000712-RN02).

## Acknowledgements

We acknowledge the gracious philanthropic funding from the Joe and Mary Moeller Foundation. This work is also supported in part by an RO1 R01CA180279 grant from the National Cancer Institute (NCI). CW also acknowledges support from the Marie-Josée Kravis Fellowship in Quantitative Biology. Funding provided by Pediatric Solid Tumors Comprehensive Data Resource Core (Grant # RP180819) and support from the Institutional Tissue Bank generated the osteosarcoma tissue data. The Flow Cytometry and Cellular Imaging Core Facility performed sorting for the osteosarcoma cells. We acknowledge the Advanced Technology Genomics Core (ATGC, Core grant: CA016672) for their support in creating single-cell libraries and sequencing the cells (NIH 1S10OD024977-01). AT was supported from AFOSR grants (FA9550-20-1-0029, FA9550-23-1-0096), Army Research Office grant (W911NF-22-1-0292), and NIH grant (R01-AG048769).

We would like to honor Dr. Allen Tannenbaum who was co-senior author on this study. Unfortunately, Allen passed away before the paper was published. We are deeply indebted to his many contributions to science as well as his positive impact on our careers.

## References

1. Jo VY, Doyle LA. Refinements in Sarcoma Classification in the Current 2013 World Health Organization Classification of Tumours of Soft Tissue and Bone. Surg Oncol Clin N Am 2016;25:621–43.

2. Thway K. Pathology of soft tissue sarcomas. Clin Oncol (R Coll Radiol) 2009;21:695–705.

3. Kansara M, Teng MW, Smyth MJ, Thomas DM. Translational biology of osteosarcoma. Nat Rev Cancer 2014;14:722–35.

4. Evola FR, Costarella L, Pavone V, et al. Biomarkers of Osteosarcoma, Chondrosarcoma, and Ewing Sarcoma. Front Pharmacol 2017;8:150.

5. Cortes-Ciriano I, Lee JJ, Xi R, et al. Comprehensive analysis of chromothripsis in 2,658 human cancers using whole-genome sequencing. Nat Genet 2020;52:331–41.

6. Beird HC, Bielack SS, Flanagan AM, et al. Osteosarcoma. Nat Rev Dis Primers 2022;8:77.

7. Grun D, Muraro MJ, Boisset JC, et al. De Novo Prediction of Stem Cell Identity using Single-Cell Transcriptome Data. Cell Stem Cell 2016;19:266–77.

8. Teschendorff AE, Enver T. Single-cell entropy for accurate estimation of differentiation potency from a cell’s transcriptome. Nat Commun 2017;8:15599.

9. Guo M, Bao EL, Wagner M, Whitsett JA, Xu Y. SLICE: determining cell differentiation and lineage based on single cell entropy. Nucleic Acids Res 2017;45:e54.

10. Gulati GS, Sikandar SS, Wesche DJ, et al. Single-cell transcriptional diversity is a hallmark of developmental potential. Science 2020;367:405–11.

11. Engler AJ, Sen S, Sweeney HL, Discher DE. Matrix elasticity directs stem cell lineage specification. Cell 2006;126:677–89.

12. Weistuch C, Murgas KA, Zhu J, et al. Cancer heterogeneity is defined by normal cellular trade-offs. bioRxiv 2023:2023.04.12.536595.

13. Truong DD, Lamhamedi-Cherradi S-E, Porter RW, et al. Dissociation protocols used for sarcoma tissues bias the transcriptome observed in single-cell and single-nucleus RNA sequencing. BMC cancer 2023;23:1–16.

14. Meyer WH, Houghton JA, Houghton PJ, Webber BL, Douglass EC, Look AT. Development and characterization of pediatric osteosarcoma xenografts. Cancer Res 1990;50:2781–5.

15. Blondel VD, Guillaume J-L, Lambiotte R, Lefebvre E. Fast unfolding of communities in large networks. Journal of statistical mechanics: theory and experiment 2008;2008:P10008.

16. Lamhamedi-Cherradi SE, Mohiuddin S, Mishra DK, et al. Transcriptional activators YAP/TAZ and AXL orchestrate dedifferentiation, cell fate, and metastasis in human osteosarcoma. Cancer Gene Ther 2021;28:1325–38.

17. Wu CC, Beird HC, Andrew Livingston J, et al. Immuno-genomic landscape of osteosarcoma. Nat Commun 2020;11:1008.

18. Wu C-L, Dicks A, Steward N, et al. Single cell transcriptomic analysis of human pluripotent stem cell chondrogenesis. Nature communications 2021;12:362.

19. Korsunsky I, Millard N, Fan J, et al. Fast, sensitive and accurate integration of single-cell data with Harmony. Nat Methods 2019;16:1289–96.

20. Pittenger MF, Mackay AM, Beck SC, et al. Multilineage potential of adult human mesenchymal stem cells. Science 1999;284:143–7.

21. Tirosh I, Izar B, Prakadan SM, et al. Dissecting the multicellular ecosystem of metastatic melanoma by single-cell RNA-seq. Science 2016;352:189–96.

22. Roson-Burgo B, Sanchez-Guijo F, Del Canizo C, De Las Rivas J. Insights into the human mesenchymal stromal/stem cell identity through integrative transcriptomic profiling. BMC Genomics 2016;17:944.

23. Kovar H, Bierbaumer L, Radic-Sarikas B. The YAP/TAZ Pathway in Osteogenesis and Bone Sarcoma Pathogenesis. Cells 2020;9.

24. Dupont S, Morsut L, Aragona M, et al. Role of YAP/TAZ in mechanotransduction. Nature 2011;474:179–83.

25. Tang Y, Feinberg T, Keller ET, Li XY, Weiss SJ. Snail/Slug binding interactions with YAP/TAZ control skeletal stem cell self-renewal and differentiation. Nat Cell Biol 2016;18:917–29.

26. Zheng X, Zeng W, Gai X, et al. Role of the Hedgehog pathway in hepatocellular carcinoma (review). Oncol Rep 2013;30:2020–6.

27. Akladios B, Reinoso VM, Cain JE, et al. Positive regulatory interactions between YAP and Hedgehog signalling in skin homeostasis and BCC development in mouse skin in vivo. PloS one 2017;12.

28. Elsafadi M, Manikandan M, Dawud R, et al. Transgelin is a TGF β-inducible gene that regulates osteoblastic and adipogenic differentiation of human skeletal stem cells through actin cytoskeleston organization. Cell death & disease 2016;7:e2321-e.

29. Zhu B, Xue F, Li G, Zhang C. CRYAB promotes osteogenic differentiation of human bone marrow stem cells via stabilizing beta-catenin and promoting the Wnt signalling. Cell Prolif 2020;53:e12709.

30. Zhou BO, Yue R, Murphy MM, Peyer JG, Morrison SJ. Leptin-receptor-expressing mesenchymal stromal cells represent the main source of bone formed by adult bone marrow. Cell Stem Cell 2014;15:154–68.

31. Sanchez C, Mazzucchelli G, Lambert C, Comblain F, DePauw E, Henrotin Y. Comparison of secretome from osteoblasts derived from sclerotic versus non-sclerotic subchondral bone in OA: A pilot study. PLoS One 2018;13:e0194591.

32. Ebert R, Benisch P, Krug M, et al. Acute phase serum amyloid A induces proinflammatory cytokines and mineralization via toll-like receptor 4 in mesenchymal stem cells. Stem Cell Res 2015;15:231–9.

33. Marie PJ. Osteoblast biology and mechanosensing. Mechanosensing Biology: Springer; 2011:105–26.

34. Jundt G, Berghauser KH, Termine JD, Schulz A. Osteonectin--a differentiation marker of bone cells. Cell Tissue Res 1987;248:409–15.

35. Abdelmagid SM, Barbe MF, Rico MC, et al. Osteoactivin, an anabolic factor that regulates osteoblast differentiation and function. Experimental cell research 2008;314:2334–51.

36. Merle B, Bouet G, Rousseau JC, Bertholon C, Garnero P. Periostin and transforming growth factor β-induced protein (TGF β I p) are both expressed by osteoblasts and osteoclasts. Cell biology international 2014;38:398–404.

37. Hauschka PV, Lian JB, Cole D, Gundberg CM. Osteocalcin and matrix Gla protein: vitamin K-dependent proteins in bone. Physiological reviews 1989;69:990–1047.

38. Smargiassi A, Bertacchini J, Checchi M, et al. WISP-2 expression induced by Teriparatide treatment affects in vitro osteoblast differentiation and improves in vivo osteogenesis. Mol Cell Endocrinol 2020:110817.

39. Gustafson B, Hammarstedt A, Hedjazifar S, Smith U. Restricted adipogenesis in hypertrophic obesity: the role of WISP2, WNT, and BMP4. Diabetes 2013;62:2997–3004.

40. Ambele MA, Dessels C, Durandt C, Pepper MS. Genome-wide analysis of gene expression during adipogenesis in human adipose-derived stromal cells reveals novel patterns of gene expression during adipocyte differentiation. Stem Cell Res 2016;16:725–34.

41. Kadota Y, Toriuchi Y, Aki Y, et al. Metallothioneins regulate the adipogenic differentiation of 3T3-L1 cells via the insulin signaling pathway. PLoS One 2017;12:e0176070.

42. Nishizuka M, Koyanagi A, Osada S, Imagawa M. Wnt4 and Wnt5a promote adipocyte differentiation. FEBS Lett 2008;582:3201–5.

43. Cheon CW, Kim DH, Kim DH, Cho YH, Kim JH. Effects of ciglitazone and troglitazone on the proliferation of human stomach cancer cells. World J Gastroenterol 2009;15:310–20.

44. Mariman EC, Wang P. Adipocyte extracellular matrix composition, dynamics and role in obesity. Cell Mol Life Sci 2010;67:1277–92.

45. Hardingham TE, Oldershaw RA, Tew SR. Cartilage, SOX9 and Notch signals in chondrogenesis. J Anat 2006;209:469–80.

46. Bougault C, Priam S, Houard X, et al. Protective role of frizzled-related protein B on matrix metalloproteinase induction in mouse chondrocytes. Arthritis Res Ther 2014;16:R137.

47. Takacs R, Vago J, Poliska S, et al. The temporal transcriptomic signature of cartilage formation. Nucleic Acids Res 2023;51:3590–617.

48. Zhang Y, Pizzute T, Pei M. A review of crosstalk between MAPK and Wnt signals and its impact on cartilage regeneration. Cell Tissue Res 2014;358:633–49.

49. Park S, Seo K, So A, et al. SOX2 has a crucial role in the lineage determination and proliferation of mesenchymal stem cells through Dickkopf-1 and c-MYC. Cell Death & Differentiation 2012;19:534–45.

50. Wang Y-X, Zhang Y-J. Nonnegative matrix factorization: A comprehensive review. IEEE Transactions on knowledge and data engineering 2012;25:1336–53.

51. Zhou Y, Yang D, Yang Q, et al. Single-cell RNA landscape of intratumoral heterogeneity and immunosuppressive microenvironment in advanced osteosarcoma. Nature communications 2020;11:6322.

52. Bentzen SM, Poulsen HS, Kaae S, et al. Prognostic factors in osteosarcomas. A regression analysis. Cancer 1988;62:194–202.

53. Chui MH, Kandel RA, Wong M, et al. Histopathologic Features of Prognostic Significance in High-Grade Osteosarcoma. Arch Pathol Lab Med 2016;140:1231–42.

54. Sun HH, Chen XY, Cui JQ, Zhou ZM, Guo KJ. Prognostic factors to survival of patients with chondroblastic osteosarcoma. Medicine (Baltimore) 2018;97:e12636.

55. Molina ER, Chim LK, Lamhamedi-Cherradi SE, et al. Correlation of nuclear pIGF-1R/IGF-1R and YAP/TAZ in a tissue microarray with outcomes in osteosarcoma patients. Oncotarget 2022;13:521–33.

56. Engler AJ, Griffin MA, Sen S, Bonnemann CG, Sweeney HL, Discher DE. Myotubes differentiate optimally on substrates with tissue-like stiffness: pathological implications for soft or stiff microenvironments. J Cell Biol 2004;166:877–87.

57. Reticker-Flynn NE, Malta DFB, Winslow MM, et al. A combinatorial extracellular matrix platform identifies cell-extracellular matrix interactions that correlate with metastasis. Nature communications 2012;3:1122.

58. Comley K, Fleck NA. A micromechanical model for the Young’s modulus of adipose tissue. International Journal of Solids and Structures 2010;47:2982–90.

59. Iivarinen JT, Korhonen RK, Julkunen P, Jurvelin JS. Experimental and computational analysis of soft tissue stiffness in forearm using a manual indentation device. Medical engineering & physics 2011;33:1245–53.

60. Zaky S, Lee K, Gao J, et al. Poly (glycerol sebacate) elastomer supports bone regeneration by its mechanical properties being closer to osteoid tissue rather than to mature bone. Acta biomaterialia 2017;54:95–106.

61. Wolf FA, Angerer P, Theis FJ. SCANPY: large-scale single-cell gene expression data analysis. Genome Biol 2018;19:15.

62. Butler A, Hoffman P, Smibert P, Papalexi E, Satija R. Integrating single-cell transcriptomic data across different conditions, technologies, and species. Nat Biotechnol 2018;36:411–20.

63. Hafemeister C, Satija R. Normalization and variance stabilization of single-cell RNA-seq data using regularized negative binomial regression. Genome Biology 2019;20:1–15.

64. Korsunsky I, Millard N, Fan J, et al. Fast, sensitive and accurate integration of single-cell data with Harmony. Nature methods 2019;16:1289–96.

65. McInnes L, Healy J, Melville J. Umap: Uniform manifold approximation and projection for dimension reduction. arXiv preprint arXiv:180203426 2018.

66. Gu Z, Eils R, Schlesner M. Complex heatmaps reveal patterns and correlations in multidimensional genomic data. Bioinformatics 2016;32:2847–9.

67. Wickham H. ggplot2–Elegant Graphics for Data Analysis.

68. Wu T, Hu E, Xu S, et al. clusterProfiler 4.0: A universal enrichment tool for interpreting omics data. Innovation (Camb) 2021;2:100141.

69. Yu G, Wang LG, Han Y, He QY. clusterProfiler: an R package for comparing biological themes among gene clusters. OMICS 2012;16:284–7.

70. Lee DD, Seung HS. Learning the parts of objects by non-negative matrix factorization. Nature 1999;401:788–91.

71. Ma X, Liu Y, Liu Y, et al. Pan-cancer genome and transcriptome analyses of 1,699 paediatric leukaemias and solid tumours. Nature 2018;555:371–6.

72. Goldman MJ, Craft B, Hastie M, et al. Visualizing and interpreting cancer genomics data via the Xena platform. Nat Biotechnol 2020;38:675–8.

73. Edgar R, Domrachev M, Lash AE. Gene Expression Omnibus: NCBI gene expression and hybridization array data repository. Nucleic acids research 2002;30:207–10.

