## Supplementary Figures for "Mapping the Single-cell Differentiation Landscape of Osteosarcoma"

^The authors declare no conflicts of interest.^

### Supplemental Figures

Supplemental Figure 1

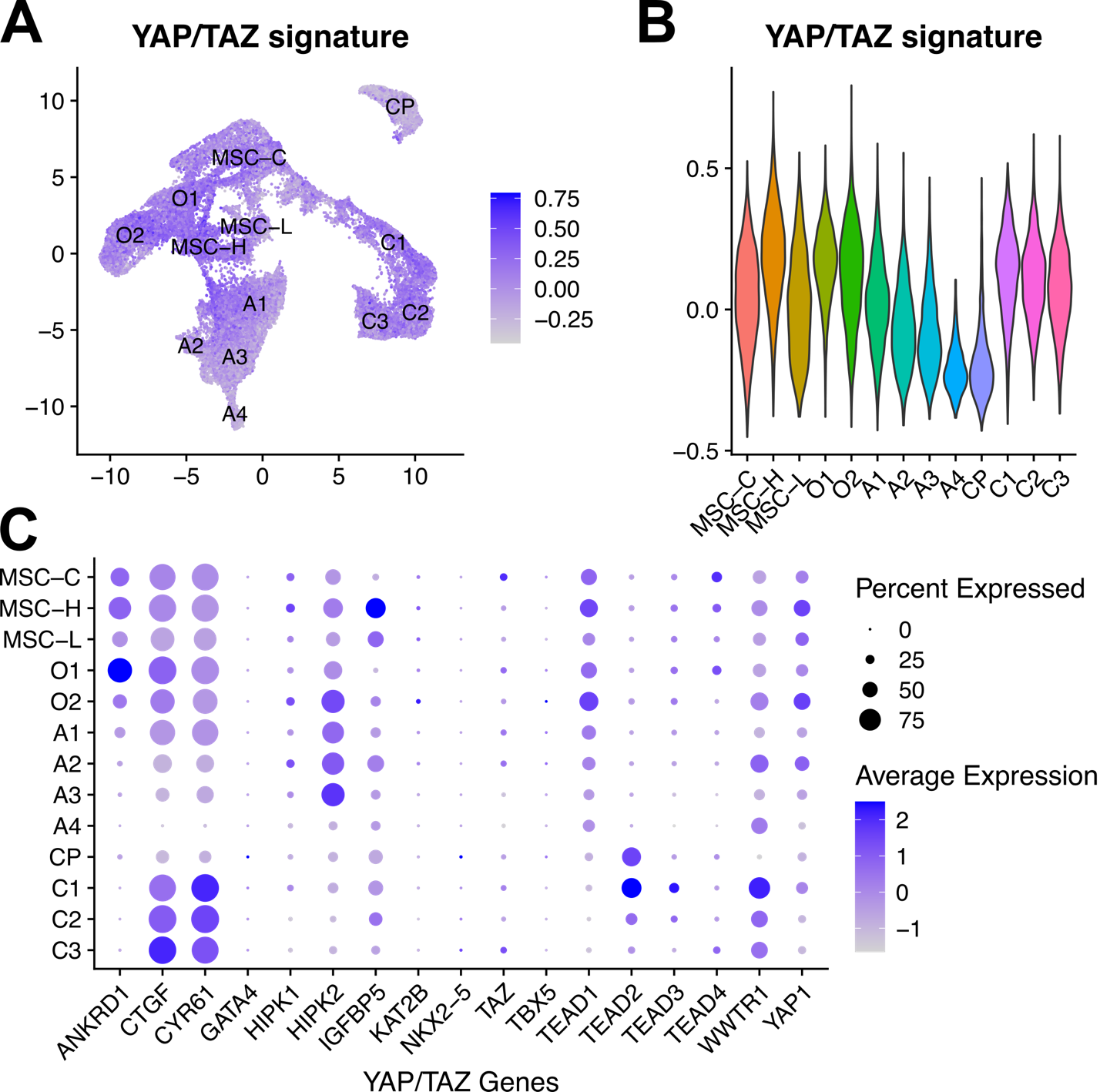

**SI Figure 1:** YAP/TAZ-related gene signatures in MTL. **A:** UMAP of YAP/TAZ signature module score. **B:** Violin plot of YAP/TAZ module score distributions across each cluster. **C:** DotPlot of individual YAP/TAZ regulated genes (and including YAP, TAZ). Color shows the average expression level of each gene, and dot size corresponds to the percentage of cells with minimal gene expression.

Supplemental Figure 2

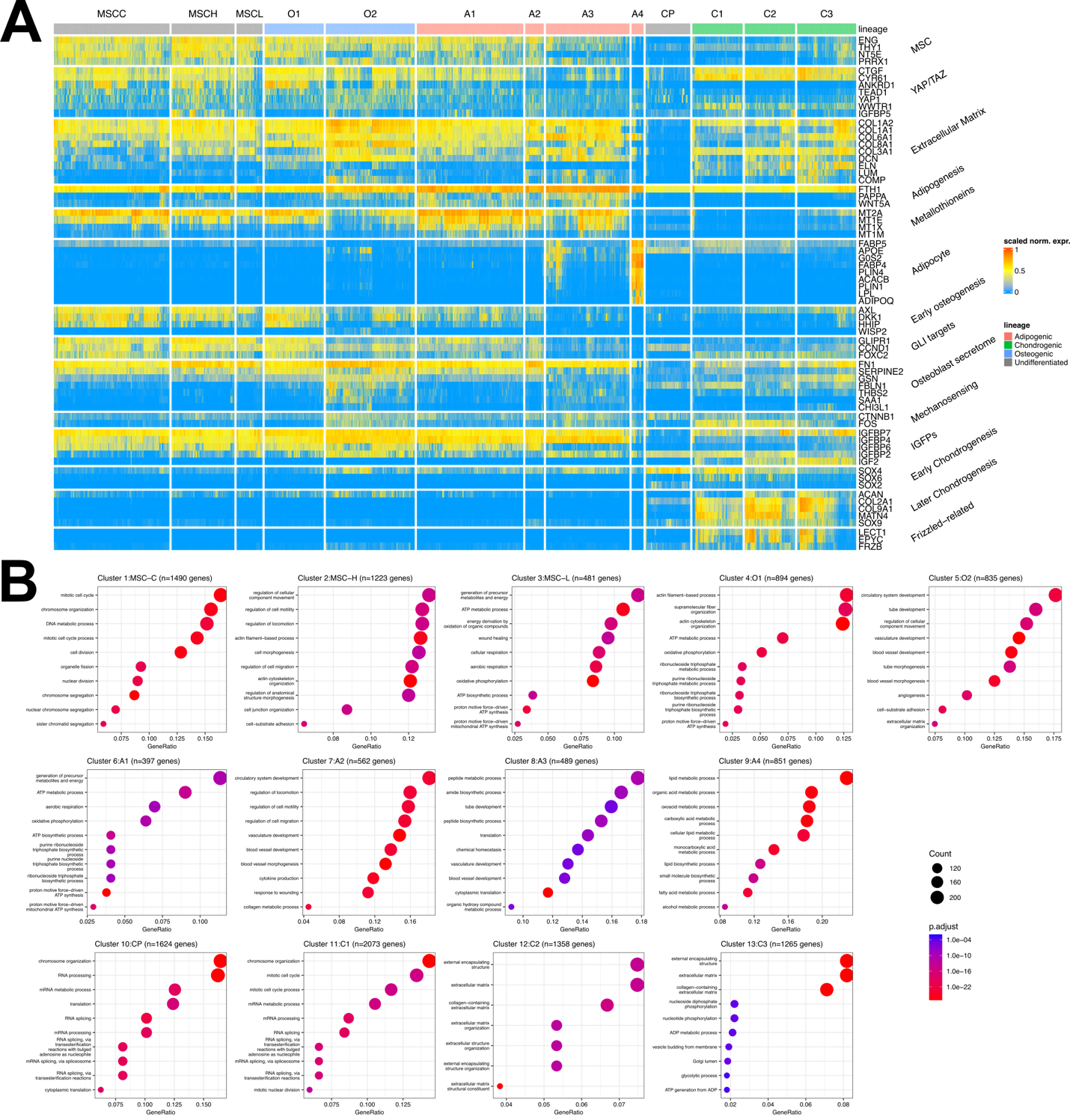

**SI Figure 2: MTL Cluster Identities are Validated by Marker Gene and Pathway Enrichment Analysis.** **A:** Heatmap of marker gene expression (SCTransform normalized data scaled to the maximum value of each gene). On the right, biological groups of genes are annotated according to known literature (see main text for references). **B:** Dotplots of Gene Ontology (GO) pathway enrichment analysis of all marker genes identified in each cluster. Here, we display the top 10 pathways, while all enrichment results are included in Supplemental Table S2. The dot color is proportional to the multiple-comparison-adjusted p-value, and the dot size corresponds to the number of marker genes in the relevant pathway.

Supplemental Figure 3

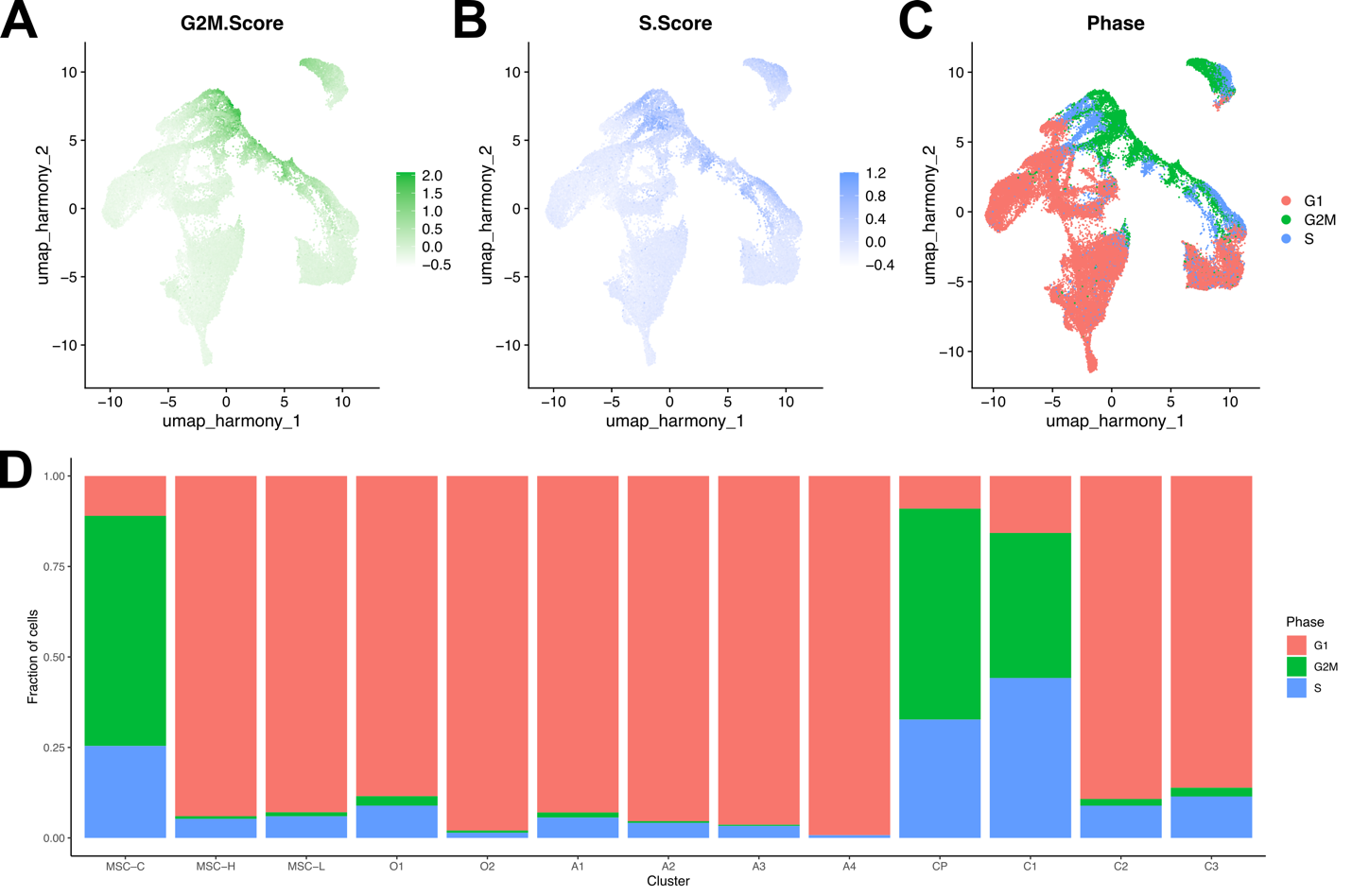
**SI Figure 3:** Cell cycle signatures in MTL. **A:** UMAP of G2/M phase module score. **B:** UMAP of S phase module score. **C:** UMAP of cell cycle phase, labeled according to G2/M and S scores. **D:** Bar plot of cell cycle phase proportions within each cluster.

Supplemental Figure 4

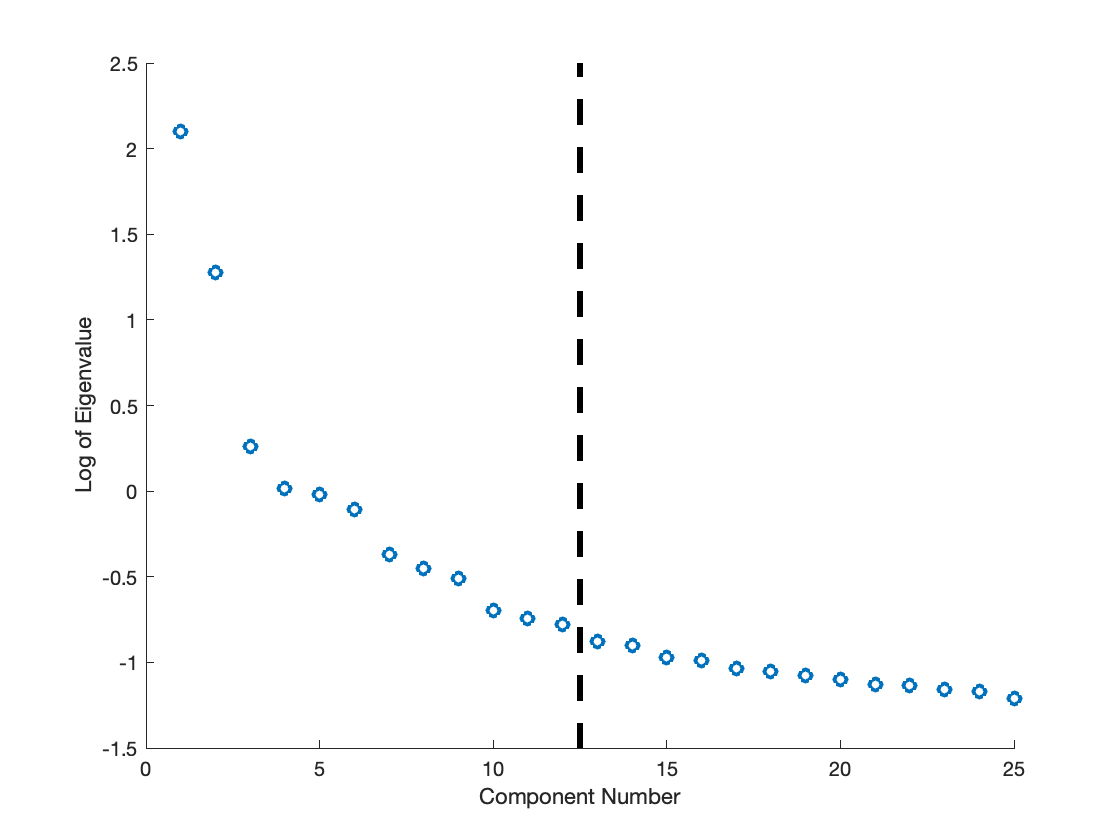

**SI Figure 4:** Scree plot of the eigenvalues of the combined MTL gene expression data.  Around the black dashed line, the eigenvalues begin to level off, indicating that 12 components are the optimal number to retain.  As further validation, additional N-NMF components did not correspond to time-dependent or lineage-specific gene expression signatures.
